# Powerful eQTL mapping through low coverage RNA sequencing

**DOI:** 10.1101/2021.08.08.455466

**Authors:** Tommer Schwarz, Toni Boltz, Kangcheng Hou, Merel Bot, Chenda Duan, Loes Olde Loohuis, Marco P. Boks, René S. Kahn, Roel A. Ophoff, Bogdan Pasaniuc

## Abstract

Mapping genetic variants that regulate gene expression (eQTL mapping) in large-scale RNA sequencing (RNA-seq) studies is often employed to understand functional consequences of regulatory variants. However, the high cost of RNA-Seq limits sample size, sequencing depth, and therefore, discovery power. In this work, we demonstrate that, given a fixed budget, eQTL discovery power can be increased by lowering the sequencing depth per sample and increasing the number of individuals sequenced in the assay. We perform RNA-Seq of whole blood tissue across 1490 individuals at low-coverage (5.9 million reads/sample) and show that the effective power is higher than that of an RNA-Seq study of 570 individuals at high-coverage (13.9 million reads/sample). Next, we leverage synthetic datasets derived from real RNA-Seq data to explore the interplay of coverage and number individuals in eQTL studies, and show that a 10-fold reduction in coverage leads to only a 2.5-fold reduction in statistical power. Our study suggests that lowering coverage while increasing the number of individuals is an effective approach to increase discovery power in RNA-Seq studies.

## BACKGROUND

The vast majority of risk loci identified in genome-wide association studies (GWAS) are difficult to interpret as they lie in noncoding regions of the genome. Variants that regulate gene expression abundance, as measured through expression quantitative trait locus (eQTL) studies, provide insightful information about the functional interpretation of GWAS signals ^1–2^. By integrating eQTL associations with GWAS, we can hope to identify target genes that are driving the GWAS signal at a locus ^3–6^. RNA sequencing (RNA-Seq) is the state-of-the-art assay for measuring gene expression in bulk tissue and is therefore the assay of choice for eQTL mapping ^7–8^. However, the high cost of RNA-Seq often limits the sample size and therefore reduces the discovery power of eQTL studies based on RNA-Seq ^2,6,9^.

Traditional RNA-Seq study design prioritizes sequencing depth per individual (targeted levels of coverage in the range of 30-50 million reads) over the number of individuals (samples) included in the study ^10–12^. However, given that high levels of coverage per individual limits the sample size of a study, this results in a loss of statistical power in eQTL mapping. Previous studies have established that the low-coverage whole genome sequencing of a larger number of individuals attains increased power of association compared to higher-coverage studies of smaller sample sizes in GWAS ^13–17^. This raises the hypothesis that, similarly as for whole genome sequencing and GWAS, lower coverage RNA-seq with a considerable increase in the number of individuals sequenced could increase power of discovery in eQTL studies ^18–21^. Currently, there is no systematic approach for determining the optimal sample size (in terms of number of sequenced individuals) and coverage to maximize eQTL discovery power.

In this work, we perform RNA-Seq in 1490 individuals at a lower coverage (average mapped read depth of 5.9 million reads/sample) and find that eQTL discovery power is better than that of an experiment with a similar budget, but with fewer individuals and higher coverage. Compared to high-coverage RNA-Seq, we find a high degree of consistency in both the gene expression as well as eQTL effects. We assess the interplay of coverage per sample and accuracy of expression estimates using synthetic RNA-Seq datasets generated by the down-sampling of real high-coverage data. Our analyses show that a sequencing experiment conducted with a target coverage of 10 million reads/sample has an average correlation per-gene of 0.40, when compared to an experiment conducted with a target coverage of 50 million reads/sample. We provide evidence to show that under a fixed budget, sequencing at lower coverage levels (< 10 million reads/sample) and increased sample size can boost the effective sample size per unit of cost compared to standard approaches of eQTL study design.

## RESULTS

### Low-coverage RNA-sequencing is accurate for eQTL mapping

To validate the utility of low-coverage RNA-sequencing, we sequenced whole blood tissue from N = 1490 unrelated individuals (**Methods**) (**Supplementary Figure 1A** and **Supplementary Figure 1B**). We target a sequencing coverage of 9.5 million reads per sample, yielding M = 5.9 million reads mapped to RefSeq genes on average (sd across samples of 1.96 million, **Supplementary Figure 2**). We refer to this dataset as the lower-coverage RNA-Seq, or the M=5.9 million reads/sample dataset. We contrast this dataset with an RNA-Seq dataset obtained with a similar budget, but with 2.4-fold higher coverage (M = 13.9 reads) across N = 570 individuals (**Supplementary Figure 1C** and **Supplementary Figure 1D**) ^22^. We refer to this as the higher-coverage whole blood RNA-Seq, or the M = 13.9 million reads/sample dataset (Table 1).

**Table 1:**
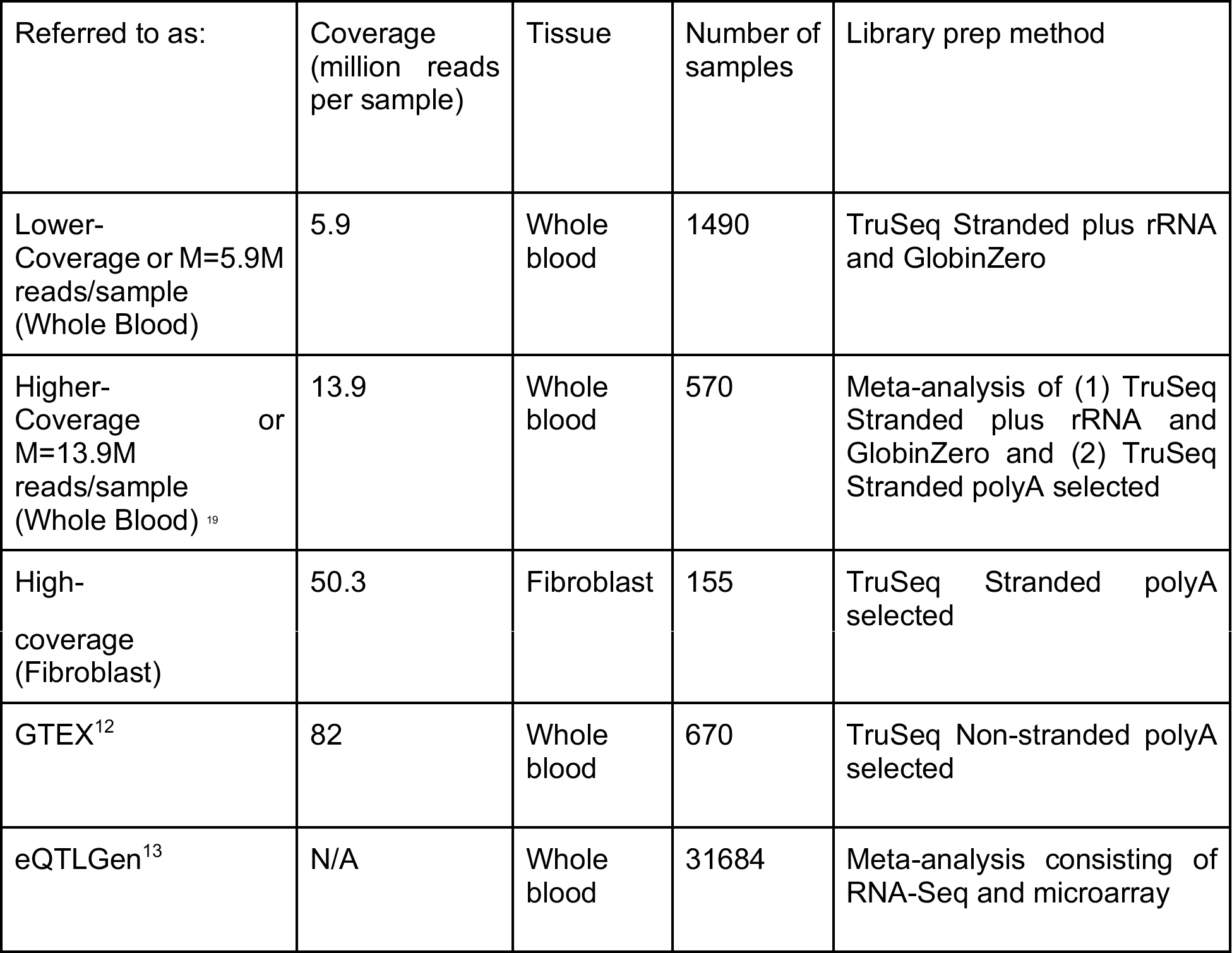
RNA-Seq datasets discussed in this paper. The coverage refers to the average number of reads that successfully map to the transcriptome, except for GTEX, which refers to the median number of total reads per sample (average mapped not available). Further description of sample overlaps among cohorts in **Supplementary Note**.

**Table 2:** Sequencing cost scenarios (Figure 3) The cost parameters corresponding to the effective sample size scenarios in Figure 3. Cost per sample reflects the cost of library prep to include an additional sample. Cost per lane reflects the cost per sequencing lane, which allows for 300 million reads.

|  | Cost per lane | Cost per sample |
| --- | --- | --- |
| Scenario 1 | \$1790 | \$87 |
| Scenario 2 | \$1790 | \$30 |
| Scenario 3 | \$1790 | \$150 |
| Scenario 4 | \$1000 | \$150 |

First, we assess the number of genes quantified in the two datasets. We observe 40459 genes with at least one mapped read on average across samples in the whole blood high-coverage dataset, and 27308 genes with at least one mapped read on average across samples in the whole blood low-coverage dataset. Notably, when restricting to protein coding genes with at least one mapped read in both the high-coverage and low-coverage datasets, we find more similar numbers between the data sets, with 18329 and 15605 genes quantified, respectively. This is likely due to the very sparse abundance of the non-protein coding genes, making them less likely to be detected in a lower coverage dataset. Indeed, we observe similar effects across the high vs low coverage datasets when assessing the genes with sufficient expression to be included in eQTL analysis (TPM > 0.1 in 20% of individuals, see **Methods**): 26566 genes (15496 protein coding genes) in high coverage data versus 19039 (13339 protein coding genes) in low coverage data. Most importantly we observe a high correlation in the abundance levels across the two data sets thus demonstrating that high and low coverage recover similar expression (R^2^= 0.91, **Figure 1A**).

**Figure 1:**
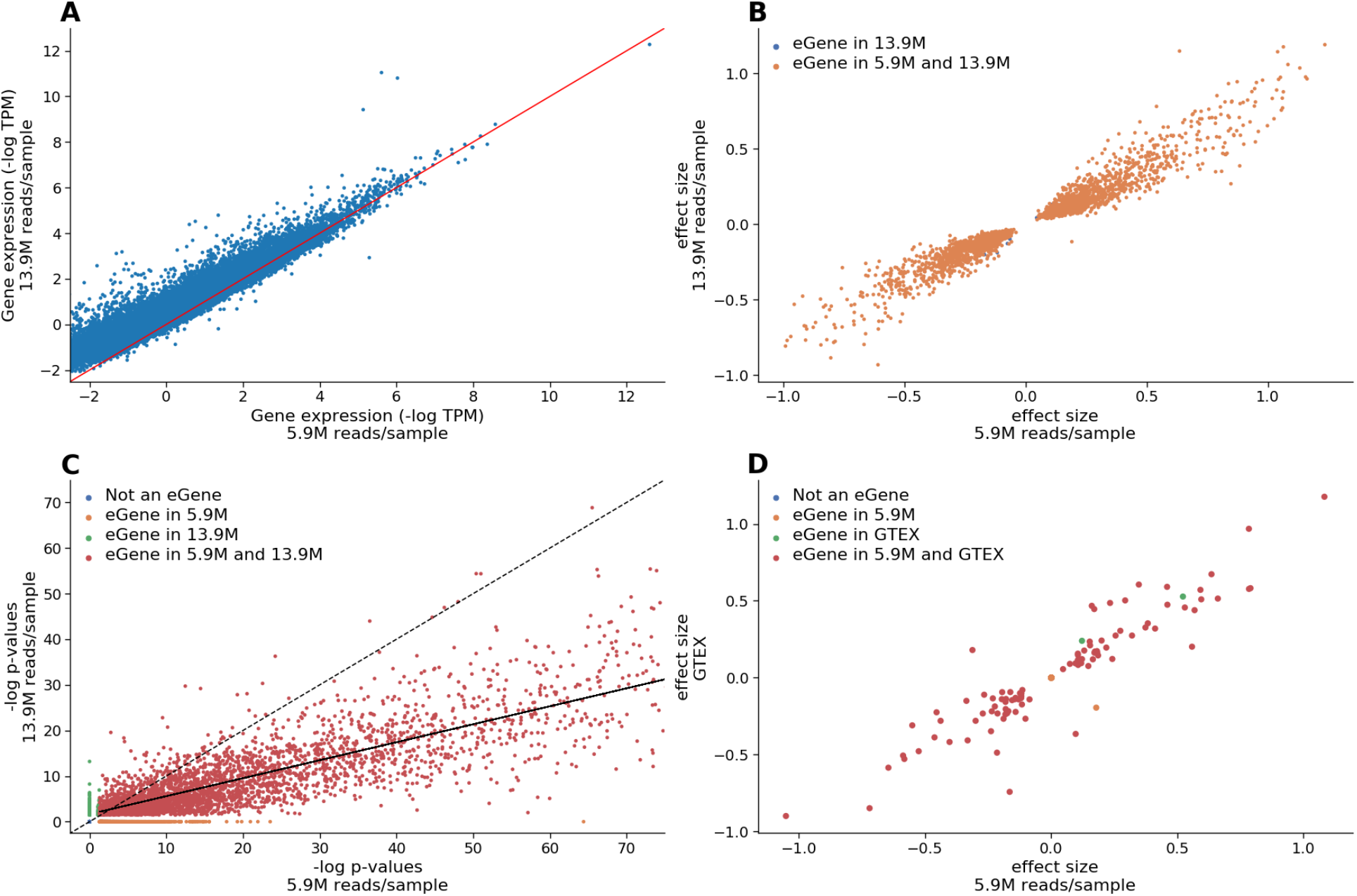
Concordance of eQTL discovery when using lower-coverage RNA-Seq vs higher-coverage RNA-Seq. **(1A):** Restricting to the 20735 genes with sufficient expression levels to be included in eQTL analysis in both the 5.9M read/sample and 13.9M read/sample dataset, comparison of the median expression (log TPM) across samples, of every gene. R^2^ = 0.91. **(1B):** In real data, scatterplot of effect sizes of most significant eQTL hits for the 2151 protein coding genes with the same eQTL hit in both eQTL analyses performed (low-coverage and high-coverage). On the x-axis, we show the effect sizes for these genes using low-coverage RNA-Seq, on the y-axis we show the effect sizes for these genes using high-coverage RNA-Seq. **(1C):** Real data p-value comparison scatterplot: In real data, scatterplot of −log p-values of most significant eQTL hit for 13950 genes included in both eQTL analyses performed (low-coverage and high-coverage). On the x-axis, we show the −log p-values for these genes using low-coverage RNA-Seq, on the y-axis we show the −log p-values for these genes using high-coverage RNA-Seq. The dotted line shows *y* = *x*, while the solid line shows the line of best fit for the 3985 protein-coding eGenes with a significant eQTL hit in both datasets. **(1D):** In real data, scatterplot of effect sizes of the most significant eQTL hit for the 140 eGenes with the same leading SNP identified in both eQTL analyses performed (lower-coverage RNA-Seq with 5.9M reads/sample and GTEX). On the x-axis, we show the effect size for these eGenes from eQTL analysis conducted using the 1490 individuals of EUR ancestry and imputed genotypes, and on the y-axis we show the effect sizes for these eGenes from eQTL analysis published by the GTEX Consortium.

Next, we investigate the power of low-coverage RNA-Seq for eQTL mapping. We conducted cis-eQTL mapping with a 1 Mb window using FastQTL, restricting to the 1490 unrelated individuals in the low-coverage RNA-Seq data (**Methods**), to identify 7587 genes (eGenes) with a significant association at FDR correction level of 5%. As expected, eQTL distribution is concentrated at transcription start sites (TSS), with 73% of eGenes TSS within 250kb of the associated SNP (eSNP). Repeating this approach using the high-coverage whole blood data in 570 individuals, we only find 5971 genes with a significant association at FDR correction level of 5%. 4969 of the 7587 eGenes found using the low-coverage data are also significant in the high-coverage data. Of these, 2151 of the eGenes are protein coding eGenes that share the same associated eSNP, and we see an extremely high level of concordance between effect sizes for these eGenes across the two datasets (R^2^ = 0.93, **Figure 1B**). This further indicates that low-coverage RNA-Seq is robust in capturing eQTL effect sizes. 1002 genes were found to be eGenes in the high-coverage eQTL analysis but not in the low-coverage analysis, with 573 (of the 1002) not passing expression levels (TPM >0.1 in 20% individuals) to be included in the low-coverage eQTL analysis; only 234 of the 573 were protein coding genes. Similar concordance is observed at the level of p-values for the associations in both datasets (**Figure 1C**). Comparing the p-values for eGenes detected in both eQTL analyses, the corresponding regression line has a slope of 0.39, consistent with lower-coverage dataset having superior statistical power to detect associations over lower-coverage dataset, and consistent with overall number of significant eQTL discoveries. We report the results from using typed SNPs in these eQTL analyses (**Methods**), but observe similar patterns when using the full set of imputed SNPs.

To further validate the performance of eQTL analysis using lower coverage RNA-Seq (coverage 5.9M, n = 1490), we compared the resulting eQTLs to the ones found by GTEx consortium in whole blood ^12^ (**Supplementary Figure 3**). Restricting to the 12247 protein coding genes with sufficient expression to be included in both studies (> 0.1 TPM in 20% of samples) we find that 4529 out of the 5957 protein coding genes (76%) with a significant association using the lower-coverage data also had a significant association in GTEx, correcting at an FDR level of 5%. Further restricting to eGenes with the same leading SNP in both of these datasets (140 genes) (**Figure 1D**), we observe a correlation (R^2^) of 0.85 between their effect sizes. Looking into the 1428 protein coding genes with a significant association in eQTL analysis using the lower-coverage RNA-Seq but not in GTEx using an FDR cutoff of 5%, we observe that 372 have significant association in GTEx using an FDR cutoff of 10%. To further ensure that these eGenes are not false positives, we compare the set of 1428 genes with eQTL analysis conducted by the eQTLGen Consortium ^23^ and find that all but 190 of these genes have been found to have a significant association in eQTLGen. This suggests that the additional associations found using lower-coverage data that are not found in GTEx are not false positives, but fall just below the significance threshold in the GTEx analysis.

Finally, we explore the impact of RNA-Seq at lower coverages for cell type expression estimation. We use Cibersort ^24^ to compare cell-type proportion estimates between the lower-coverage data and higher-coverage data (**Methods**). We find that the median estimated cell type proportions are conserved across both datasets, suggesting that deconvolution of cell type specific signal from gene expression profiles of whole blood samples is not impacted when coverage is reduced by half (**Supplementary Figure 7**).

### Impact of RNA-seq coverage on eQTL power

Having demonstrated the accuracy of low-coverage RNA-Seq in eQTL mapping in real data, we next focused on exploring the interplay of number of individuals and coverage for optimizing power for discovery. As simulating RNA-Seq data is challenging ^25–26^, we down-sample reads from higher-coverage RNA-Seq data to create synthetic datasets at various coverages (**Methods**). We observe that with just a fraction of the reads, it is still possible to estimate gene expression (**Figure 2A**). For example, using just 10% of the data (5.0 million reads/sample) retains a per gene R^2^ of 0.40, on average. The results from our analyses using these synthetic lower-coverage RNA-Seq datasets suggest that under simplified settings of no per-sample library preparation cost, the statistical power in an association study can be increased up to fourfold by spending more resources on increasing sample size and fewer resources on increasing coverage. In practice, increasing the number of samples in an RNA-Seq study leads to increased library preparation costs, making the increase in obtainable statistical association power less obvious.

**Figure 2:**
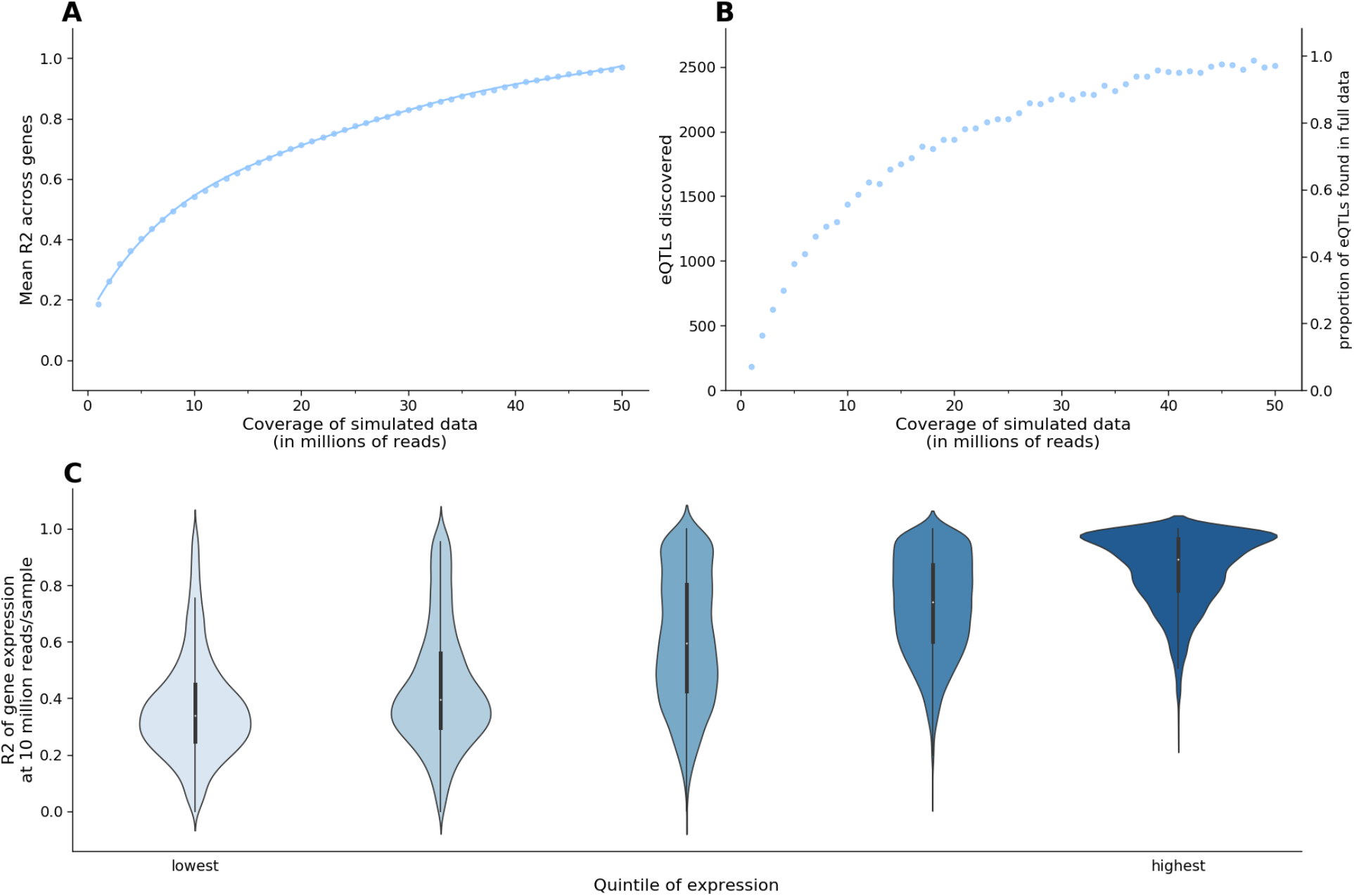
Synthetic lower-coverage RNA-Seq captures expression signal. **(2A):** On the x-axis, we show the level of simulated coverage, and on the y-axis we show the mean Pearson correlation of every gene. We calculate this value by finding the R^2^ values for the TPM values of each of 45,910 genes across 155 samples between the high coverage data (average of 50 million reads per sample) and the simulated data, and reporting the mean R^2^ value per gene. **(2B):** For a fixed number of individuals, absolute number and percentage of eGenes captured at 5% FDR, for synthetic RNA-Seq at varying levels of coverage. **(2C):** Gene expression accuracy as a function of relative gene expression observed in actual RNA-Seq data with 50 million reads/sample. 23,540 genes (with average expression < 0.1 TPM) are divided into five ascending quintiles of expression based on their average expression in 155 samples.

It has been established that statistical power in association studies is a function of sample size, phenotype measurement accuracy, and genotype measurement accuracy ^13,19,29^. This means that the power of a study with sample size N and estimated gene expression is approximately the same as the power of a study with sample size N, using the true gene expression measurements (**Methods**). In this scenario, R^2^ is the correlation between the true expression and the expression estimates. We therefore report the squared correlation (R^2^) between synthetic datasets at various coverages and the full data at an average of 50 million reads/sample (which is assumed to be the true gene expression). While these results show the mean R^2^ for all genes obtained under one synthetic dataset (one draw) per coverage level, we find that the synthetic datasets are consistent across multiple draws at the same coverage level (**Supplementary Figure 4A**) and each show similar correlations with the ground truth gene expression (**Supplementary Figure 4B**).

Next, we quantified how well lower-coverage RNA-Seq can be used to detect eGenes ^30^. We explore the number of genes with significant associations after FDR correction at 5% under various levels of simulated coverage (**Figure 2B**). Using synthetic data, as the number of reads per sample decreases, we find that many eGenes are still detectable. For example, at 10 million reads per sample, just 20% of the full coverage, 60% of the eGenes are still detected. In the context of eQTL studies, synthetic RNA-Seq supports the idea that sequencing at lower coverages over a higher number of individuals is a promising approach to boosting statistical power.

Finally, we explore the estimation accuracy in the synthetic data as a function of relative gene expression abundance, since less abundant genes may not be captured altogether at lower sequencing coverages. We stratify genes into five groups based on their relative expression in the full dataset (M=50.3 million reads/sample) and report the R^2^ for genes in each of these groups in synthetic data (**Figure 2C**). We observe that in the synthetic RNA-Seq dataset at 10 million reads/sample, we capture expression of highly expressed genes better than lower expressed genes. Specifically, for genes in the lowest through the highest quintiles of relative gene abundance, we find the average correlation (R^2^) to the ground truth of expression to be 0.36, 0.44, 0.61, 0.73, 0.86, respectively. We observe the same effect for synthetic datasets at coverages of 1 million reads/sample and 25 million reads/sample (**Supplementary Figure 5A** and **Supplementary Figure 5B**). These results suggest that the ability to achieve similar power in eQTL analysis studies will differ per gene, and is a function of relative expression. We further investigate the properties of genes with quantification accuracy influenced by coverage levels of sequencing and find that that protein coding genes are more accurately quantified at lower coverage levels (**Supplementary Figure 6A**). Conversely, the number of transcripts per gene, gene length, and GC content do not appear to be factors that broadly influence the gene quantification accuracy when sequencing coverage is reduced (**Supplementary Figure 6B**, **Supplementary Figure 6C**, and **Supplementary Figure 6D**).

### Optimal association power for eQTLs is attained at lower coverage with a larger number of samples

In the context of reducing experimental costs, we explored the trade-off between the number of samples sequenced and the average coverage per sample. We evaluated the expected effective sample size obtained with lower coverage per sample and compared this with a conventional approach of 50 million reads/sample. We down-sample reads (as described in Section 1 and **Methods**) from a high-coverage RNA-Seq experiment derived from Fibroblast tissue in order to create lower-coverage RNA-Seq synthetic data. This is done to match actual low coverage sequencing as closely as possible. To evaluate the relationship between cost, coverage, and sample size, we use the following equation to model the budget: 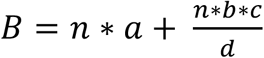

Where *B* is the cost/budget (in US dollars), *a* is the library preparation cost per sample, *b* is the target coverage of each sample (in millions of reads), *c* is the cost per lane (which contains *d* million reads), and *d* is the number of reads per sequencing lane (in millions). We compute the effective sample size of an eQTL study as a function of average coverage, which determines the number of samples sequenced under a fixed budget (**Figure 3A**). As an example, at a fixed budget of $300,000, the highest effective sample size is achieved by sequencing 2045 individuals using 10 million reads per sample, which leads to a corresponding effective sample size of 1107. An experiment achieving the sample effective sample size, using 50 million reads per sample, would cost $426,564 (N = 1107, R^2^ = 1.0). Therefore, by lowering the coverage of each sample and increasing sample size, we achieve the same effective sample size at just 70.3% of the cost.

**Figure 3:**
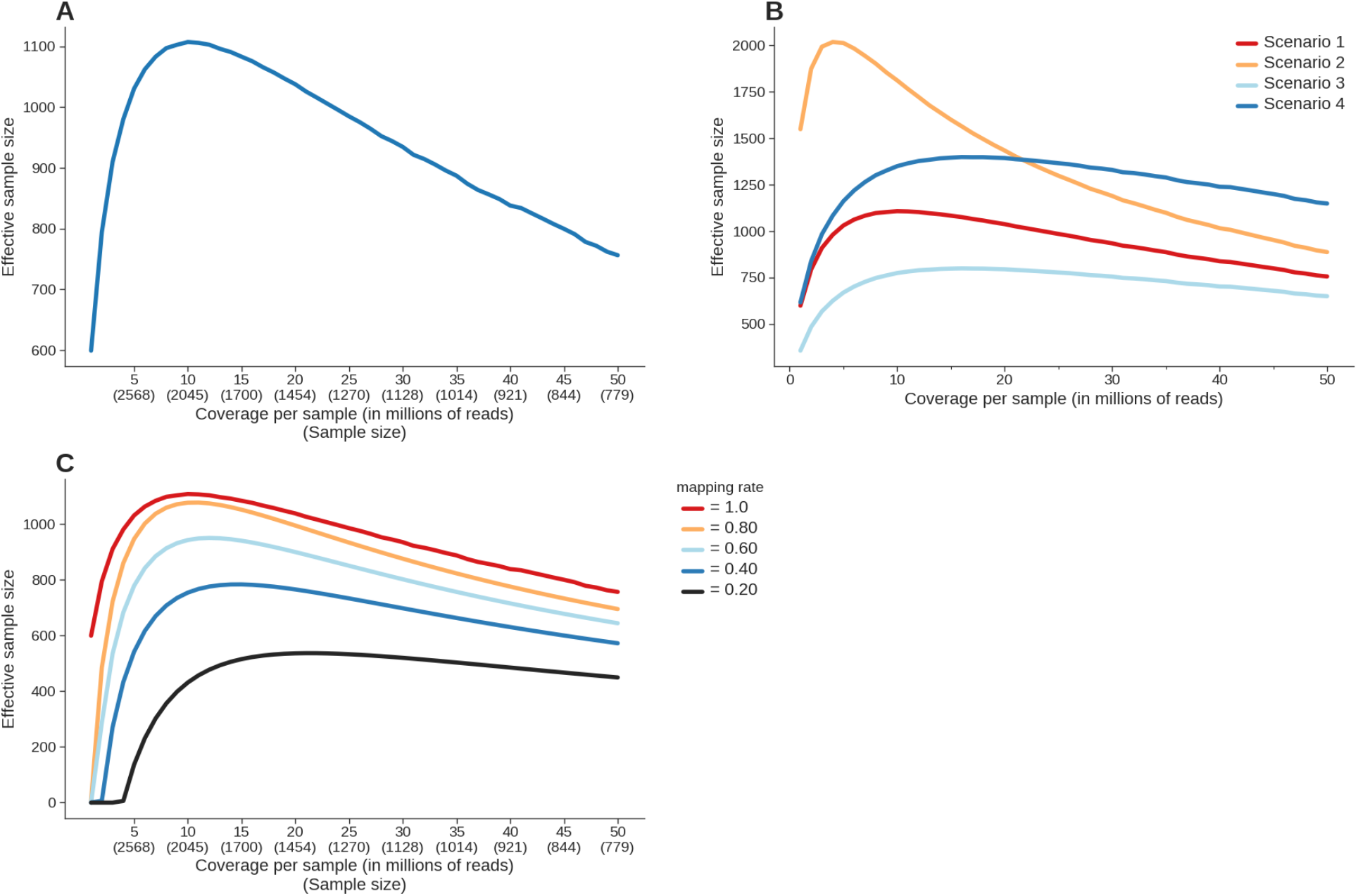
Effective sample size under various budget parameters. **(3A):** Effective sample size in RNA-Seq under a fixed budget ($300,000) as a function of the number of samples and the resulting coverage. Cost assumptions: $87 per library prep per sample, $1790 per lane of sequencing (300 million reads). **(3B):** Effective sample size in RNA-Seq under a fixed budget ($300,000) as a function of the number of samples and the resulting coverage. Cost assumptions vary and are reflected in Table 2. **(3C):** Effective sample size under a fixed budget ($300,000) as a function of the number of samples and the results coverage. A global mapping rate parameter is used to simulate actual experimental conditions (Methods).

In practice, it is common to observe a considerable discrepancy between the target number of reads in an experiment and the number of reads that successfully map to genes. This can be attributed to different library prep techniques, quality of samples, or tissue type. To show how mapping rate can influence the effective sample size of an experiment, we model effective sample size with varying levels of mapping rates (**Methods**). As expected, we observe that as the mapping rate increases, there is a corresponding increase in effective sample size (**Figure 3C**). We provide a webtool as a practical approach for selecting cost-effective designs for maximizing eQTL power: https://tomschwarz.shinyapps.io/RNASeqCoverageCalculator/.

With a budget of ~$300k and an expected mapping rate of 0.60 (chosen based on mapping rate of similar experiments using TruSeq Stranded plus rRNA and GlobinZero in whole blood tissue), we see the maximum effective sample size would be achieved at a target coverage of 7 million reads per sample, including 2227 individuals in the study. We estimate that achieving the same effective sample size using data with 50 million reads per sample would cost ~$500k (N = 1328), or 1.78x the cost of sequencing 2227 individuals at a coverage of 7 million reads/sample. To explore other cost scenarios we provide a webtool we created a webtool where one can enter budget, costs, and other details about the experiment, in order to see how to achieve optimal effective sample size (https://tomschwarz.shinyapps.io/RNASeqCoverageCalculator/).

## DISCUSSION

In this work, we generate RNA-Seq data at a lower coverage than typically used in eQTL studies (5.9M reads/sample) and demonstrate how this approach boosts effective sample size per unit cost in an association study. To further validate this approach, we use synthetic RNA-Seq data to show that the optimal level of coverage in an RNA-Seq project for the purpose of identifying eQTL associations is lower than is commonly practiced ^10–12^. Based on our findings, we recommend increasing sample size while lowering sequencing depth per sample in order to achieve optimal statistical power in association studies.

We conclude with some notes, caveats, and future directions. First, synthetic RNA-Seq via downsampling reads is potentially limited in several ways. These synthetic datasets of lower coverage RNA-Seq are created by uniformly sampling from real RNA-Seq data with an average of 50 million reads mapped per sample. However, in practice, it is possible that sequencing biases are not captured by uniform sampling due to the different experimental setup compared to the dataset from which we sample ^27^. Additionally, these synthetic datasets are based on data obtained from fibroblast tissue with different transcriptomic profiles from whole blood, potentially influencing the sequencing depth required to detect associations with gene expression. Finally, this approach is optimized for eQTL discovery. Other mechanisms that are detected using RNA-Seq, such as RNA splicing, have different mechanisms and will likely have different optimal coverages for detection. The fact that we identify different sets of eGenes depending on which gene expression measurements we consider (GTEX vs eQTLGen vs lower-coverage RNA-Seq), shows that we need to increase cohort sizes in order to fully understand the connection between genetics and gene expression in blood. Furthermore, the results in **Figure 3A** (figure showing effective sample size at various coverages) indicate that even including 1490 individuals under this fixed budget is not enough to achieve the optimal effective sample size. Current approaches are not sufficient to understand the full landscape of eQTLs in whole blood tissue, even while only considering a single genetic ancestry group. We compare the eGenes identified by GTEx, eQTLGen, and the lower-coverage RNA-Seq (**Supplementary Figure 8**) and find that no single study is sufficient in capturing all of the associations in whole blood. As observed in GWAS, much larger sample sizes including far more ancestral diversity in these samples will enable discovery of novel associations in transcriptomics. Including non-European populations and considering the temporal aspect of gene expression will help us gain a more complete understanding of the blood transcriptome landscape in the entire population.

## CONCLUSIONS

In summary, we show that reducing coverage and increasing the number of samples in an eQTL study is a valid approach for increasing effective sample size of the association study. We use both real and synthetic RNA-Seq data to confirm the benefit of increased sample sizes in eQTL studies. This approach can be applied to any dataset for which genotypes are available and will help scientists optimize resources when measuring gene expression for the purpose of integration with genetics. We provide an online tool to assist with improved design of eQTL studies at https://tomschwarz.shinyapps.io/RNASeqCoverageCalculator/.

## METHODS

### Cohort Description

The samples included are from a study with individuals ascertained for bipolar disorder (BP). The cohort consists of 916 individuals with BP, 358 controls, and 216 relatives of the individuals with BP.

### Connection between effect size and R^2^

If *g* is the genotype at the SNP that we are testing for associations, and is the effect size of that SNP when regressing on the true gene expression, y, and is the effect size of that SNP when regressing on the estimated gene expression, 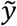. The relationship between *y* and 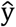 is as follows that 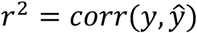. It follows that the estimates of effect size for a SNP on the true gene expression, 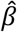, are related to the estimate of effect size for a SNP on the estimated gene expression, 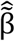 as 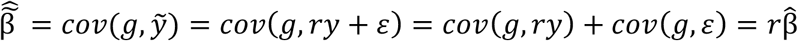 where *ε* is a random variable with mean 0 and variance 1. The association test statistics at low-coverage is *x_ground_* = *Ncor*^2^(*g, y*) thus implying that the association statistic at low coverage is 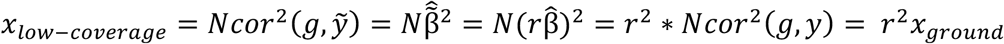

### Budget model

We modeled the cost of a large-scale bulk RNA-Seq experiment based on parameters from two different library prep techniques: (1) TruSeq Stranded plus rRNA and GlobinZero and (2) TruSeq Stranded polyA selected, both from the UCLA Neuroscience Genomics core. Cost, or *B*, is a function of the following: *a*, the library preparation cost per sample, *b*, which is the target coverage of each sample (in millions of reads). *c*, the cost per lane (which contains *d* million reads), and *d* is the number of reads per sequencing lane (in millions). Altogether, we model the budget as 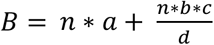

### Genotyping pipeline

Genotypes for the low-coverage whole blood samples were obtained from the following platforms: OmniExpressExome (N = 810), PSYCH (N = 523), and COEX (N = 163). Given that the SNP-genotype data for both the fibroblast and whole blood samples came from numerous studies using various genotyping platforms (including GSA, Illumina550, OmniExpress Exome, COEX, and PsychChip) the number of overlapping SNPs across all platforms was < 80k, prompting us to perform imputation separately for each genotyping platform. Genotypes were first filtered for Hardy-Weinberg equilibrium p value < 1.0e-6 for controls and p value < 1.0e-10 for cases, with minor allele frequency (MAF) > 0.01, leaving 148613 typed SNPs.

Genotypes were imputed using the 1000 Genomes Project phase 3 reference panel ^33^ by chromosome using RICOPILI v.1 ^34^ separately per genotyping platform, then subsequently merged. Imputation quality was assessed by filtering variants where genotype probability > 0.8 and INFO score > 0.1, resulting in 2289732 autosomal SNPs. We restricted to only autosomal due to sex chromosome dosage, as commonly done ^12^.

### Synthetic low coverage RNA-Seq

We use high-coverage RNA-Seq (average of 50 million reads/sample, TruSeq Stranded plus rRNA and GlobinZero library prep) from a set of 152 cell lines derived from human fibroblast cells. We assume this to be the ground truth of gene expression. We used seqtk (https://github.com/lh3/seqtk) to randomly sample reads at various coverages, uniformly. We performed five iterations of downsampling at each level of coverage in order to account for potential variability in the sampling and sequencing errors.

### RNA-Seq processing pipeline

We used FASTQC to visually inspect the read quality from the lower-coverage whole blood RNA-Seq (5.9M reads/sample) and the higher-coverage fibroblast RNA-Seq (13.9M reads/sample). We then used kallisto to pseudoalign reads to the GRCh37 transcriptome and quantify estimates for transcript expression. We aggregated transcript counts using scripts from the GTEX consortium (https://github.com/broadinstitute/gtex-pipeline) ^12^.

### cis-eQTL mapping

Excluding related individuals (pi_hat > 0.2) from the analysis, we perform cis-eQTL analysis mapping using FastQTL ^30^, using a defined window of 1 Mb both up and downstream of every gene’s TSS, for sufficiently expressed genes. We run the eQTL analysis in permutation pass mode (1000 permutations, and perform multiple testing corrections using the q-value FDR procedure, correcting at 5% unless otherwise specified. We then restrict our associations to the top (or leading) SNP per eGene.

### Cell type proportion estimation

We estimate the proportion of cell types of both the lower-coverage and higher-coverage bulk whole blood RNA-seq datasets using CIBERSORTx ^35^ with batch correction applied and LM22 signature matrix as the reference gene expression profile. The LM22 signature matrix uses 547 genes to distinguish between 22 human hematopoietic cell phenotypes.

### R^2^ adjustment

To account for the variability in mapping rate across different library prep techniques ^37^ and different tissue types, we look at the mean R^2^ at the expected coverage, which is calculated as *expected coverage = target coverage * estimated mapping rate*. Using mean R^2^ values from comparing lower-coverage synthetic RNA-Seq to higher-coverage RNA-Seq real data, we fit a log curve to estimate the adjusted mean 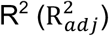 at the expected coverage.

### Effective Sample Size

Under a fixed-budget setting, we calculate effective sample size (*N_eff_*) for a given coverage using the adjusted mean 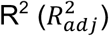 and the number of samples included at a given coverage level (N) 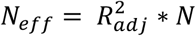

## DECLARATIONS

### Ethics approval and consent to participate

The authors assert that all procedures contributing to this work comply with the ethical standards of the relevant national and institutional committees of human experimentation and with the Helsinki Declaration of 1975, as revised in 2008.

### Consent for publication

Not applicable.

### Availability of data and materials

Gene expression data will be made available upon publication

### Author Contributions

T.S., B.P., and R.O. initialized the study. B.P. and R.O. directed and supervised the project. R.O., R.K., and M.P.B. collected samples. M.B. prepared samples for sequencing. Bioinformatics analysis was conducted by T.S., T.B., K.H., C.D., and L.O.L.. The first draft of the manuscript was drafted by T.S. and all authors contributed to editing, revisions, and approval.

## Acknowledgements

We thank the study subjects for their willingness to provide specimens and clinical data. We thank Yi Ding, Kathryn Burch, Ruthie Johnson, Arjun Bhattacharya, and Malika Freund for meaningful discussion in helping make this work possible.

## SUPPLEMENTAL INFORMATION

**Supplementary Figure 1:** *Distribution of ancestry among samples*. **(S1A)** MDS plot of 2000 samples in 5.9M read/sample cohort. **(S1B)** Distribution of ancestry among sample in 5.9M read/sample cohort. **(S1C)** MDS plot of 759 samples in 13.9M read/sample cohort: Genotype PC1 and PC2 are projected onto PCs from 1000 Genomes Project. **(S1D)** Distribution of ancestry among sample in 13.9M read/sample cohort.

**Supplementary Figure 2**: *Number of pseudoaligned reads per sample*. **(S2A)** Number of pseudoaligned reads per sample in low-coverage RNA-Seq. **(S2B)** Number of pseudoaligned reads per sample in high-coverage RNA-Seq.

**Supplementary Figure 3:** *Real data p-value comparison scatterplot with GTEX*

**Supplementary Figure 4:** *Variability in correlations in synthetic data*. **(S4A)** Scatterplot of log TPM of different draws in synthetic data. **(S4B)** Distribution of correlations observed between synthetic lower-coverage RNA-Seq and ground truth.

**Supplementary Figure 5:** *Using synthetic data, how well do we capture expression as a function of average expression in a given gene*. **(S5A)** Correlation as a function of relative expression, at 25 million reads/sample. (**S5B)** Correlation as a function of relative expression, at 1 million reads/sample.

**Supplementary Figure 6:** *Using synthetic data, how well do we capture expression in different gene categories*. (**S6A)** Using synthetic data, how well do we capture expression as a function of whether a gene is protein coding or not. **(S6B)** Using synthetic data, how well do we capture expression as a function of number of isoforms in a given gene. **(S6C)** Using synthetic RNA-Seq, how well do we capture expression as a function of gene length in a given gene Gene expression estimation accuracy simulated at 10 million reads/sample as a function of relative gene length. **(S6D)** Using synthetic RNA-Seq, how well do we capture expression as a function of GC content.

**Supplementary Figure 7:** *Estimation of cell-type proportions*.

**Supplementary Figure 8:** *Overlap of significant eGenes using RNA-Seq from three different datasets*

