## Supplementary Note for "Powerful eQTL mapping through low coverage RNA sequencing"

Notes about overlap in datasets:

- The samples in the low-coverage whole blood and high-coverage whole blood datasets are completely disjoint – no individuals overlap here.
- 97 individuals have RNA-seq data in both the high-coverage fibroblast dataset and low-coverage whole blood dataset
- 41 individuals have data in the high-coverage fibroblast dataset and the high-coverage whole blood dataset
- In total, 138 individuals overlap between the high-coverage fibroblast RNA-Seq samples and whole blood RNA-Seq samples (low-coverage and high-coverage)
