## Supplementary Figures for "Powerful eQTL mapping through low coverage RNA sequencing"

S1A:

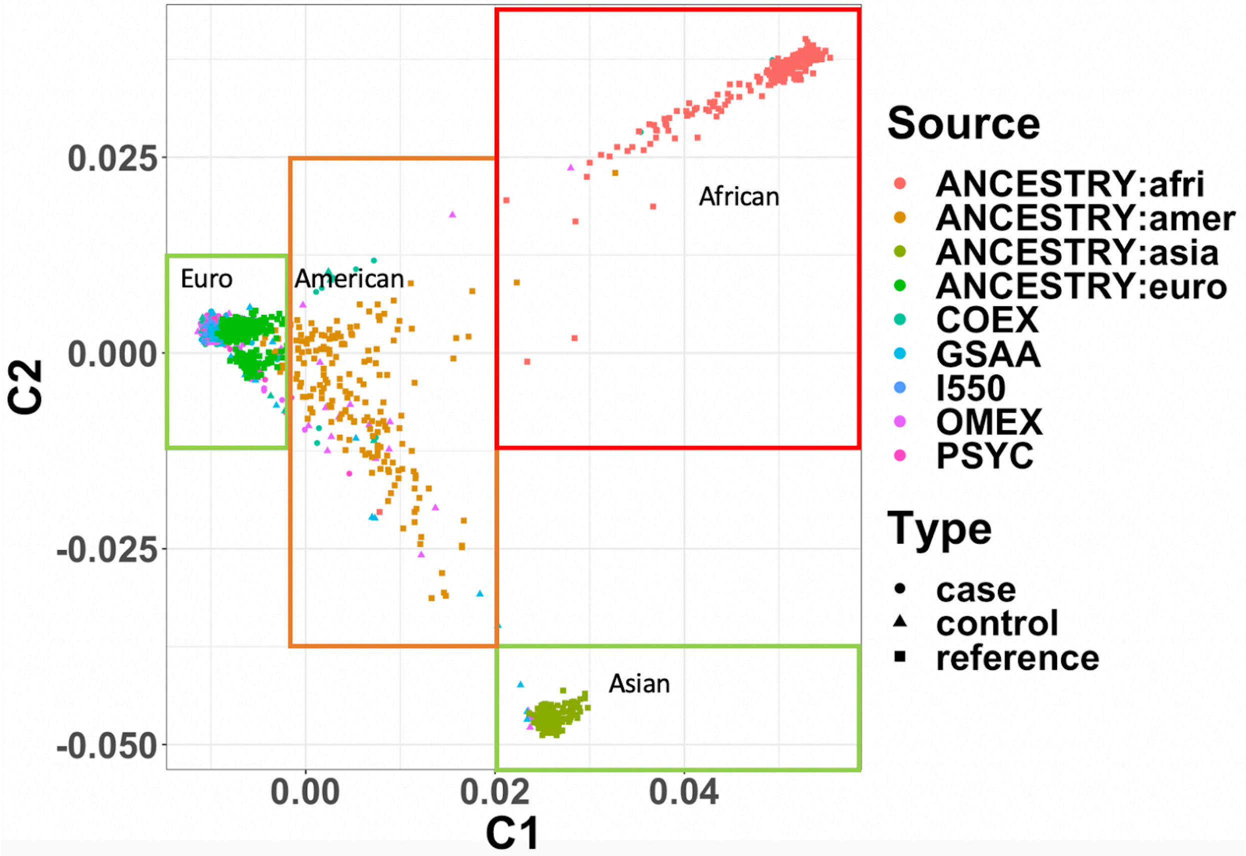


S1B:

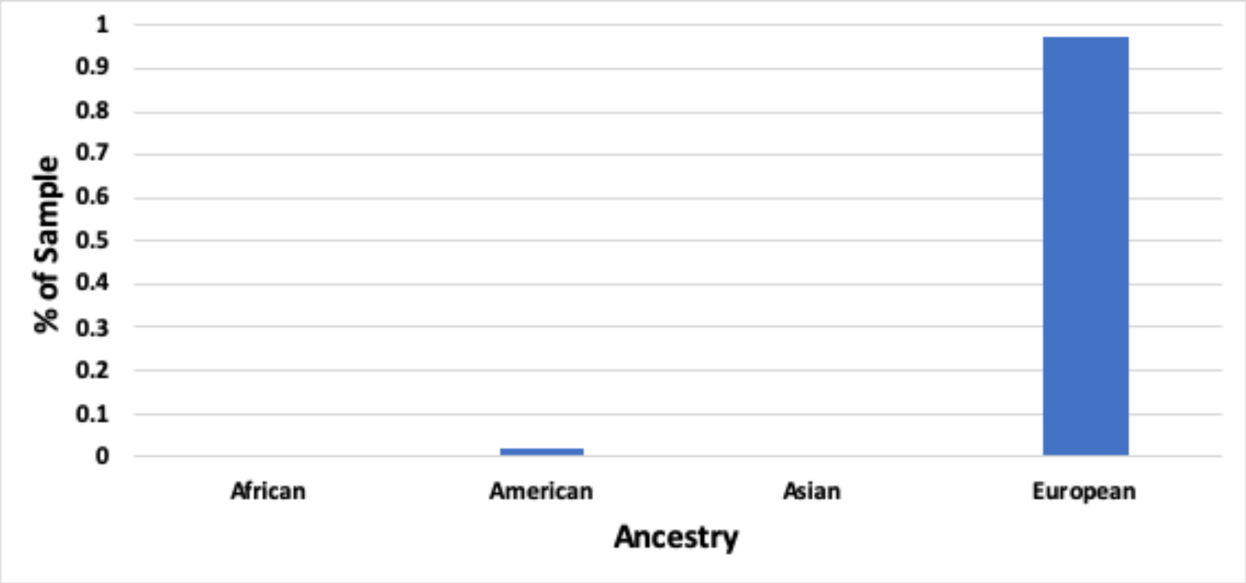

**S1C**:

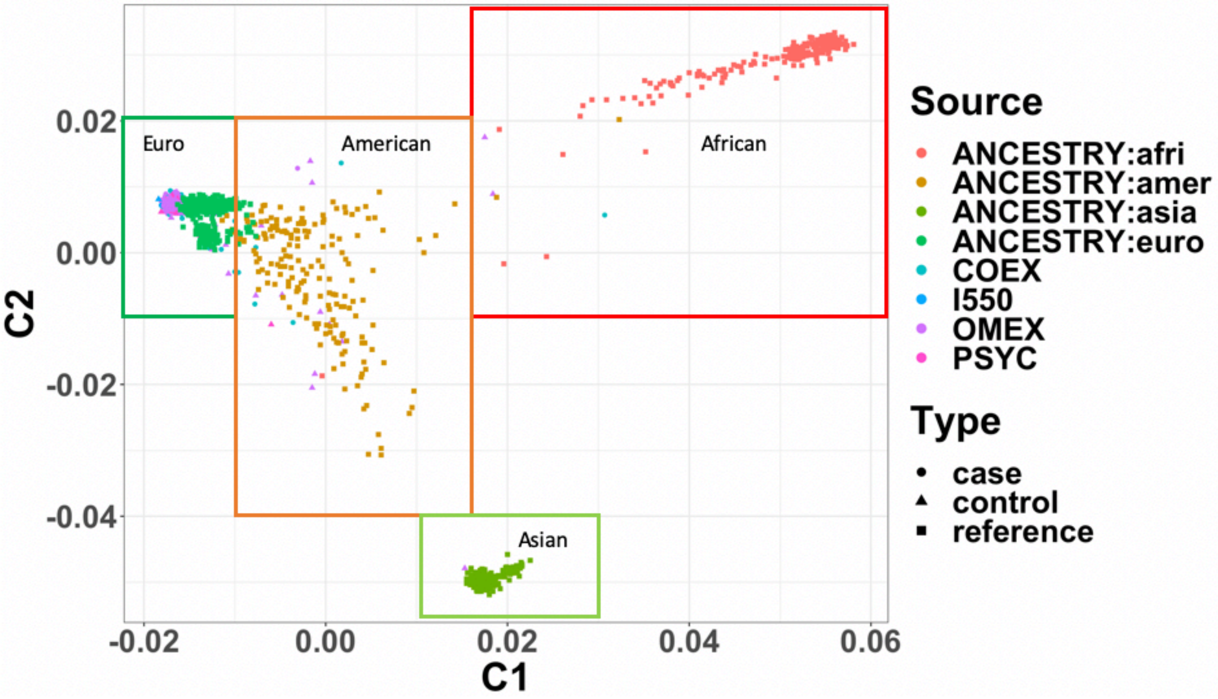


**S1D**:

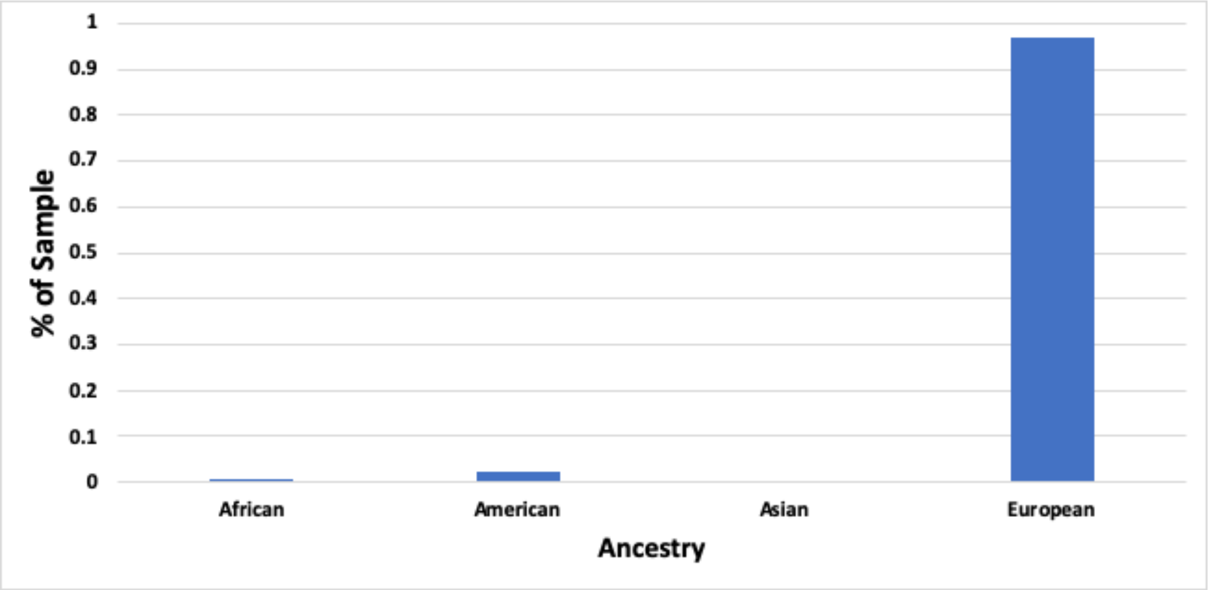


**Supplementary Figure 1:** *Distribution of ancestry among samples.* **(S1A)** Genotype PC1 and PC2 are projected onto PCs from 1000 Genomes Project. Points labeled with "ANCESTRY" are from 1000 Genomes Project, remaining points designate the specific genotyping platform used in our cohort. Boxes are drawn around the centers to show where samples from the n = 2000 / 5.9M reads/sample cohort lie. **(S1B)** A barplot showing the distribution of ancestry observed in the n = 2000 / 5.9M reads/sample cohort, according to the MDS plot. Note that only the 1963 samples that pass genotype QC thresholds are included here. Exact numbers of samples per ancestry group are: African - 4, American - 34, Asian - 9, European – 1916. **(S1C)**: Genotype PC1 and

PC2 are projected onto PCs from 1000 Genomes Project. Points labeled with “ANCESTRY” are from 1000 Genomes Project, remaining points designate the specific genotyping platform used in our cohort. Boxes are drawn around the centers to show where samples from the n = 759 / 13.9M reads/sample cohort lie. **(S1D)** A barplot showing the distribution of ancestry observed in the n = 759 / 13.9M reads/sample cohort, according to the MDS plot. Exact numbers of samples per ancestry group are: African - 4, American - 19, Asian - 1, European – 735.

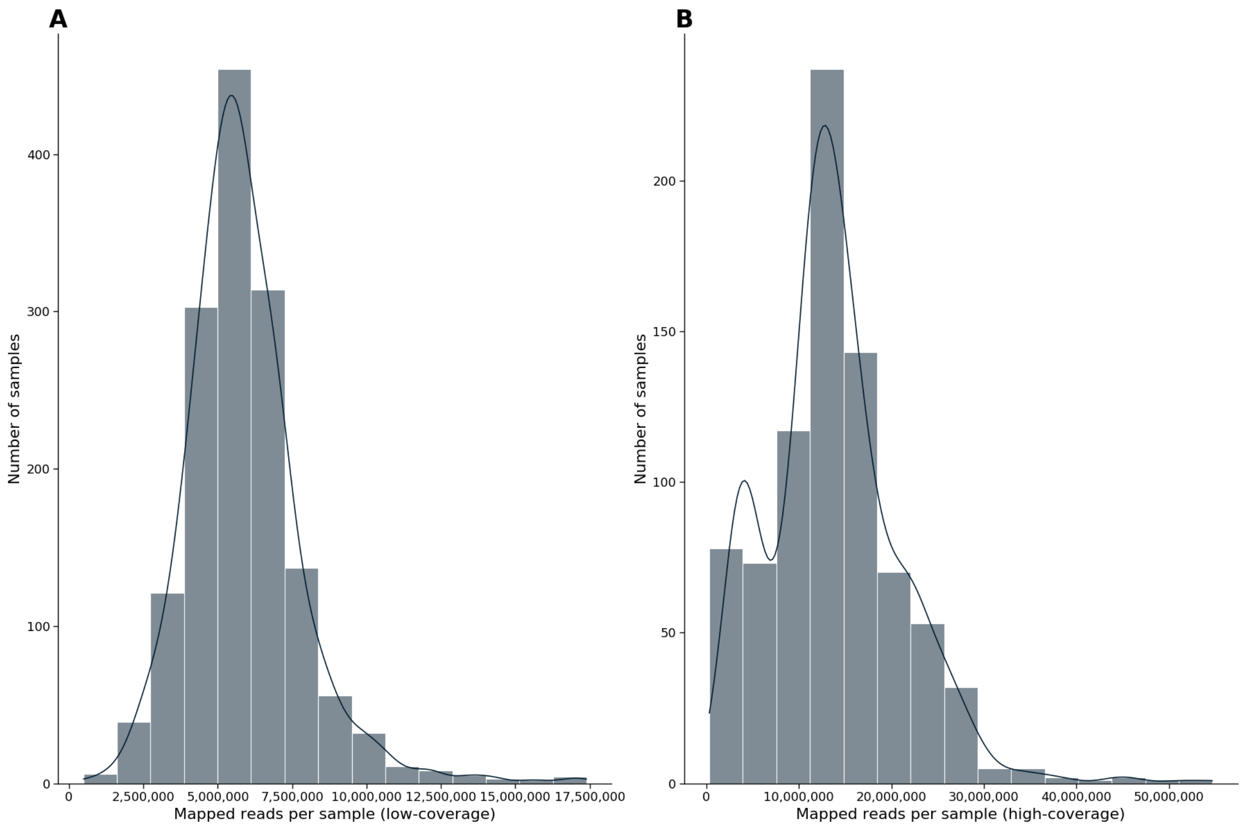


**Supplementary Figure 2:** *Number of pseudoaligned reads per sample for low-coverage and high-coverage experiments*. **(S2A)** In real data, a histogram showing the number of reads mapped to genes (or in kallisto terms: number of reads for which transcriptome successfully mapped), per sample. **(S2B)** In real data, a histogram showing the number of reads mapped to genes (or in kallisto terms: number of reads for which transcriptome successfully mapped), per sample.

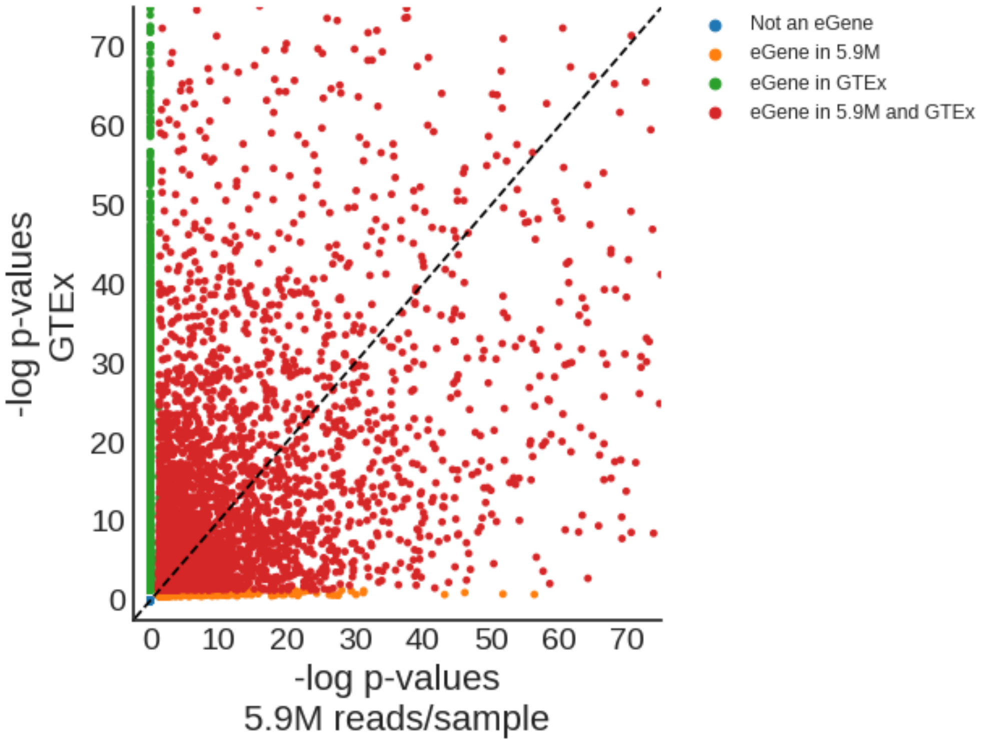


**Supplementary Figure 3:** *Real data p-value comparison scatterplot with GTEX*. **(S3A)** Using the 12,496 protein-coding genes included both in GTEX and the low-coverage datasets, on the x-axis, we show the -log p-values for leading SNP eQTL associations in the low-coverage dataset. On the y-axis, we show the -log p-values for leading SNP eQTL associations in the GTEX dataset.

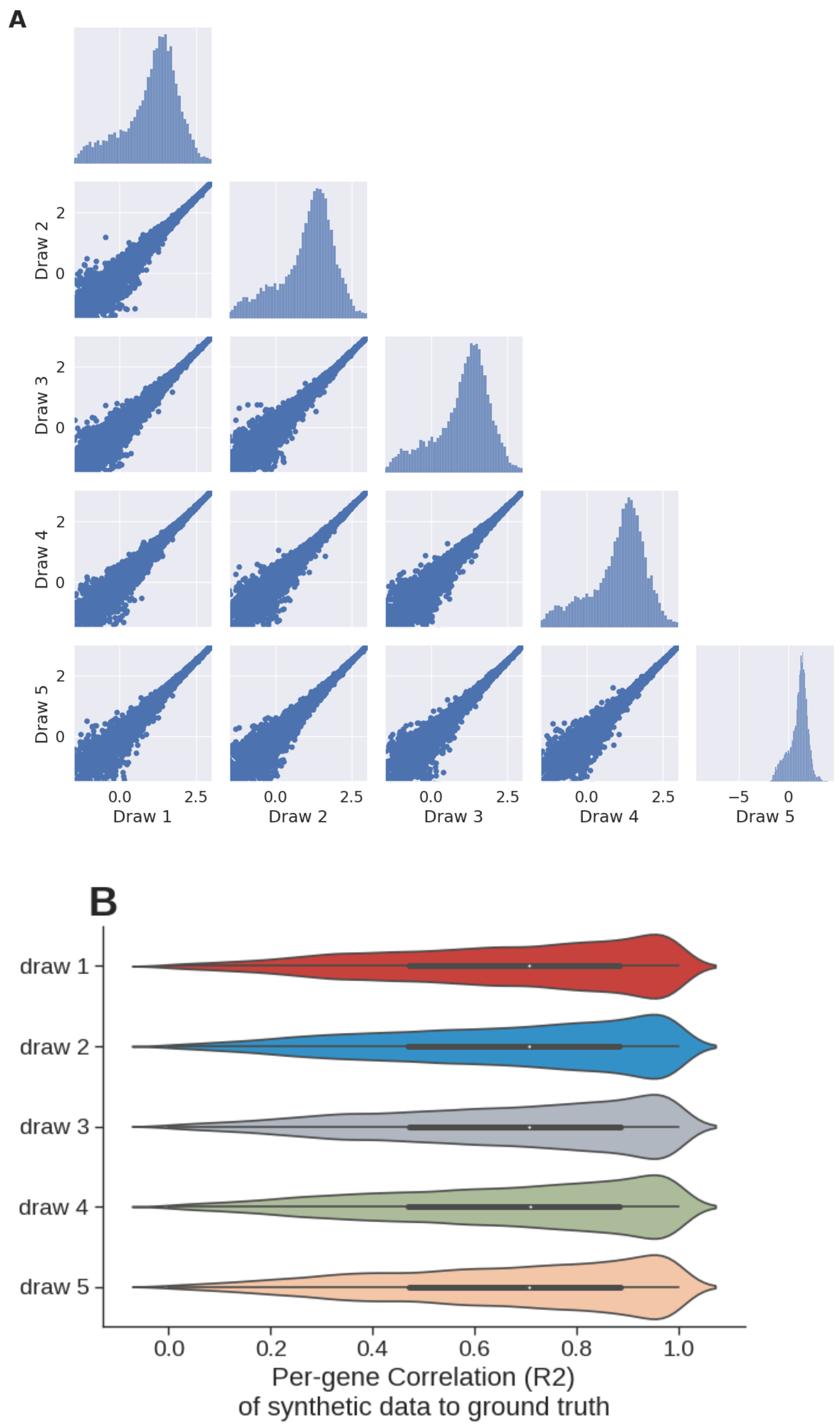


**Supplementary Figure 4:** V*ariability in correlations in synthetic data*. **(S4A)** For synthetic data corresponding to one sample, a comparison of estimated log TPM values between five different uniform sampling draws at 10 million reads/sample, for 14,948 protein-coding genes. **(S4B)** For 14,948 protein-coding genes estimated across five different uniform sampling draws at 10 million reads/sample, we compare the distribution of correlation ($R^2$) between the estimated expression of the samples and the ground truth gene expression.

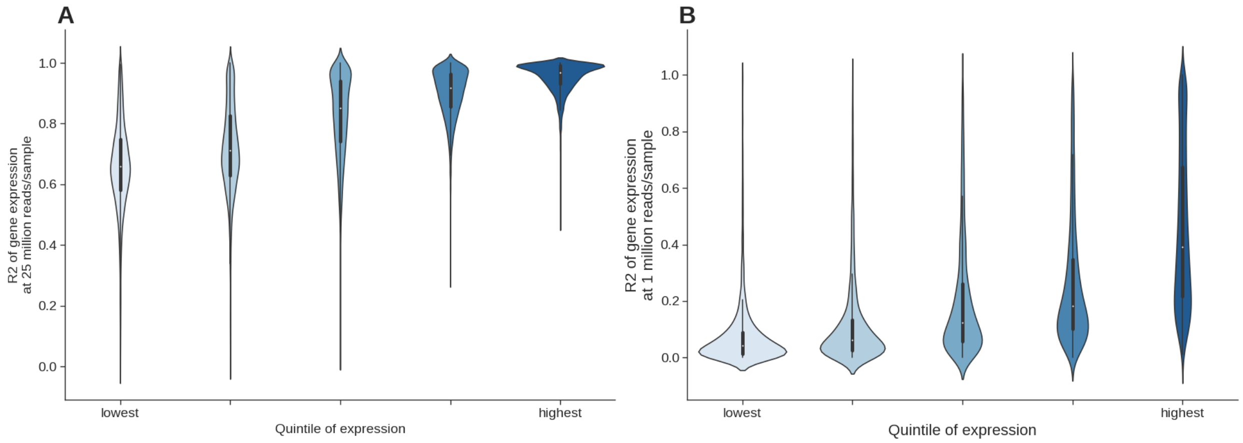


**Supplementary Figure 5:** *Using synthetic data, how well do we capture expression as a function of average expression in a given gene*. **(S5A)** Gene expression accuracy using data simulated with 1 million reads/sample, as a function of relative gene expression observed in actual RNA-Seq data with 50 million reads/sample. 23,043 genes (with average expression < 0.1 TPM) are divided into five ascending quintiles of expression based on their average expression in 155 samples. **(S5B)** Gene expression accuracy using data simulated with 1 million reads/sample, as a function of relative gene expression observed in actual RNA-Seq data with 50 million reads/sample. 23,043 genes (with average expression < 0.1 TPM) are divided into five ascending quintiles of expression based on their average expression in 155 samples.

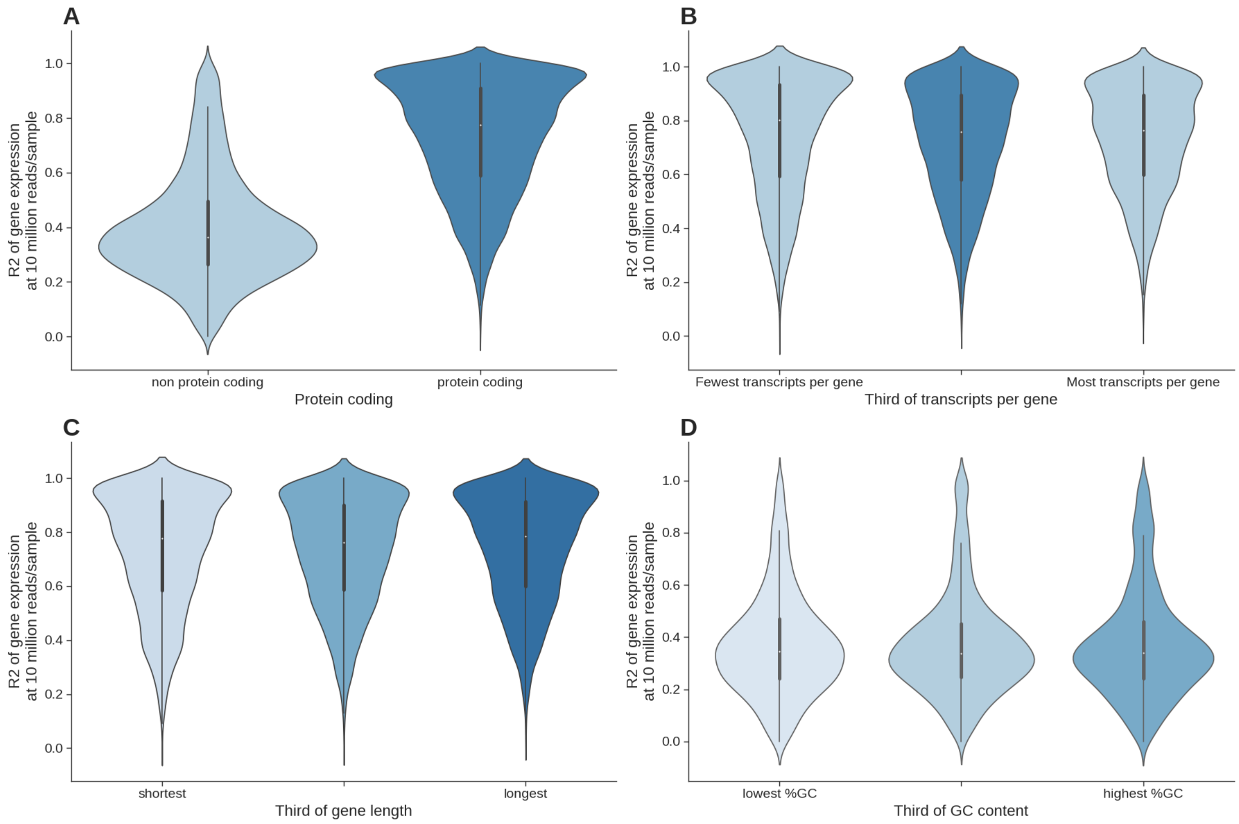


**Supplementary Figure 6:** *Using synthetic data, how well do we capture expression in different gene categories*. **(S6A)** Gene expression estimation accuracy simulated at 10 million reads/sample as a function of whether a gene codes for a protein. 24,093 genes (with average expression < 0.1 TPM) are divided into two groups. **(S6B)** Gene expression estimation accuracy simulated at 10 million reads/sample as a function of how many transcripts each gene has. 23,540 genes (with average expression < 0.1 TPM) are divided into three ascending groups based on the number of transcripts contained in each gene. **(S6C)** Gene expression estimation accuracy simulated at 10 million reads/sample as a function of relative gene length. 14,484 genes (with average expression < 0.1 TPM, protein coding) are divided into three groups based on the length of each gene. **(S6D)** Gene expression accuracy as a function of relative GC content. 5,771 genes (with average expression < 0.1 TPM and GC content reported) are divided into three groups based on the length of each gene.

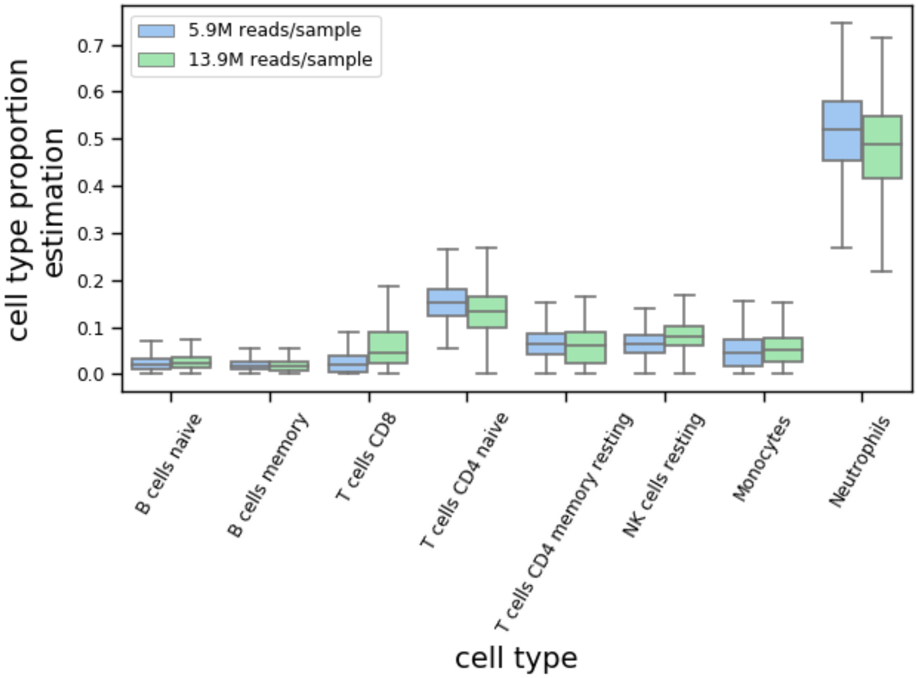


**Supplementary Figure 7:** *Estimation of cell-type proportions.* **(S7)** In real data, a comparison of estimated cell type proportions from Cibersortx between lower-coverage (5.9M reads/sample) and higher-coverage (13.9M reads/sample) RNA-Seq data for the eight most common cell types in whole blood tissue.

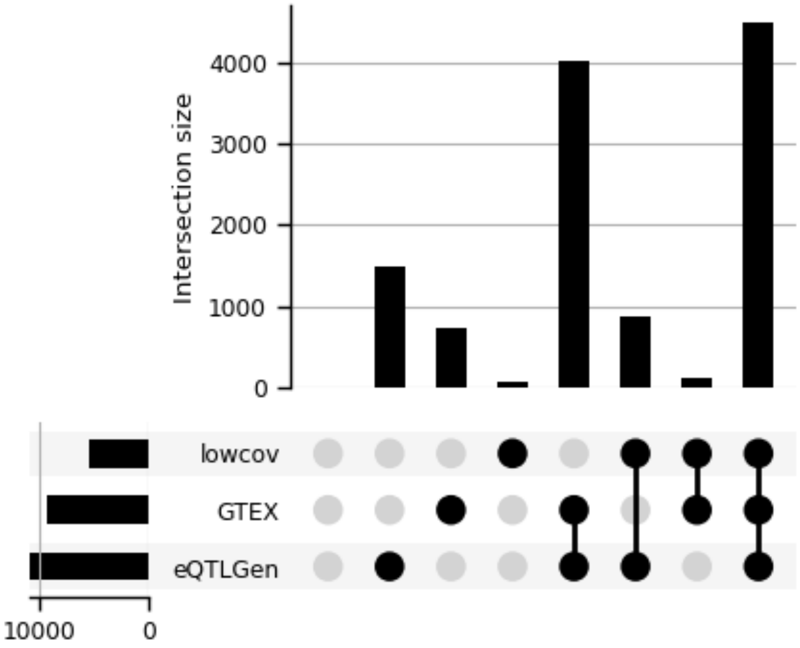


**Supplementary Figure 8:** *Overlap of significant eGenes using RNA-Seq from three different datasets*. **(S8)** Comparing number of genes with significant associations between three datasets: (1) Lower-coverage RNA-Seq (5.9M reads/sample on average, across 1,496 individuals), (2) GTEX (83M reads/sample on average, across 670 samples), (3) eQTLGen (31,684 individuals, mix of RNA-Seq and MicroArray assays used).
